# The impact of evolutionary processes in shaping the genetics of complex traits in East Asia and Europe: a specific contribution from Denisovan and Neanderthal introgression

**DOI:** 10.1101/2021.08.12.456138

**Authors:** Dora Koller, Frank R Wendt, Gita A Pathak, Antonella De Lillo, Flavio De Angelis, Brenda Cabrera-Mendoza, Serena Tucci, Renato Polimanti

## Abstract

Evidence of how human evolution shaped the polygenicity of human traits and diseases has been extensively studied in populations of European descent. However, limited information is currently available about its impact on other ancestry groups. Here, we investigated how different evolutionary processes affected the common variant heritability of traits and diseases in East Asians. Leveraging genome-wide association statistics from the Biobank Japan (up to 158,284 participants), we assessed natural selection (negative and positive), archaic introgression from Neanderthal and Denisova, and several genomic functional categories with respect to the heritability of physiological and pathological conditions. Similar to reports in European descent populations, the heritability estimates for East Asian traits were ubiquitously enriched for negative selection annotations (false discovery rate, FDR q<0.05). Enrichment of Denisovan introgression was identified in coronary artery disease (1.69-fold enrichment, p=0.003). We followed up these enrichments by conducting a phenome-wide association study (PheWAS) of Denisovan and Neanderthal alleles in participants of six ancestral backgrounds from the UK Biobank. In East Asians, Denisovan-inherited alleles were associated with 22 phenotypes, including metabolic, immunological, cardiovascular, endocrine, and dermatological traits. The strongest association was observed for the Denisovan-inherited locus rs59185462 with rheumatoid arthritis (beta=0.82, p=1.91×10^−105^). In summary, our study provides the first evidence regarding the impact of evolutionary processes on the genetics of complex traits in worldwide populations, highlighting the specific contribution of Denisovan introgression in East Asian populations.

## Introduction

Genetic variation across worldwide populations reflects the widespread impact of human evolutionary history, including processes related to natural selection and demographic history ^1^. Large-scale genome-wide association studies (GWAS) are disentangling the complex genetic architecture of human traits and diseases, providing insights into the molecular and cellular mechanisms at the basis of physiological and pathological conditions ^2–5^. Leveraging genome-wide data from these studies, it is possible to investigate whether the SNP-based heritability (SNP-*h*^*2*^, i.e., the proportion of phenotypic variance explained by additive effects of common genetic variation) of human phenotypes is enriched for specific genomic features ^6^. Genomic features related to natural selection are enriched for loci associated with complex traits ^1,7–9^. In particular, background selection (i.e., the selective removal of deleterious alleles across the genome) appears to play a primary role in shaping the highly polygenic architecture of human traits and diseases ^1,7–9^. On the other hand, genic and loss-of-function intolerant (LOF) regions are signatures of negative selection ^10^, and regions with high CpG content are positively correlated with genic content ^11^. Selection usually occurs in the form of negative selection, however, positive selection, a measure for adaptive evolution was also detected in complex traits previously ^12–14^. Genomic signatures of positive selection include soft selective sweep and hard sweep ^15^. In particular, genomic regions including enhancers present an accelerated evolutionary rate, a signature of positive selection ^13^ alike abnormally long haplotypes ^16^ and extended haplotype homozygosity ^17^.

Introgression from Neanderthals and Denisovans, the only archaic humans sequenced to date, also contributes to the genetic pool of modern populations ^18,19^ and consequently to the human phenotypic spectrum ^20,21^. Signatures of introgression in several traits, (e.g., hair and skin traits and immunity ^21–25^, neoplasms and metabolic traits ^25–27^, and male sterility ^23,28^) were identified from Neanderthals and Denisovans. The genomic segments of anatomically modern humans inherited from the admixture events with extinct human species are hypothesized to have contributed to the adaptation processes of worldwide populations that occurred during the colonization of landmasses ^23–25,28,29^. In populations of European descent (EUR), a phenome-wide association study of Neanderthal-introgressed alleles showed a wide range of associations with physiological conditions related to the immune system, skin pigmentation, and metabolic pathways, and with pathological outcomes such as depression, actinic keratosis, hypercoagulation, and tobacco use ^20^. Due to the well-known disparities of ancestry representation in biomedical research, the information currently available regarding the role of human evolutionary history in shaping the genetic architecture of traits and diseases is mostly for EUR individuals. A few studies were performed in Pacific ^30^, East Asian ^31^, Tibetan ^32^ and Island South East ^33^ populations, however, none of these studies considered genome-wide and locus-level evidence. Yet, gaps in knowledge exist on the role of certain evolutionary processes that did not occur in EUR populations. This major gap has important implications for the characterization of the history of human populations and its phenotypic consequences on individuals of diverse ancestral backgrounds.

The present study aimed to investigate how genomic elements explain part of the polygenic inheritance of human diseases and traits across different ancestry groups. In addition to baseline genomic features, we focused on understanding the impact of evolutionary processes. Leveraging data generated from large-scale GWAS conducted in the Biobank Japan (BBJ) ^34,35^, we analyzed the genetic background of individuals of East Asian descent (EAS). Populations in East Asia present an evolutionary history that is only partially shared with EUR populations. For instance, earlier studies found that on average, an EAS individual carries a higher percentage of Neanderthal genome DNA than an EUR individual (1.4% and 1.1%, respectively) ^23^. However, recent studies report that this could be due to the underestimation of archaic sequence in EUR as a result of migration back to Africa ^36^ and due to the early gene flow from modern humans to Neanderthals ^37^. EAS populations also show evidence of introgression from Denisovans ^25,28^, which is almost null in EUR populations ^18^. Accordingly, we explored how functional elements and evolutionary processes (e.g., natural selection and archaic introgression) contributed to the genetics of complex traits in EAS populations compared to EUR populations. We also conducted a phenome-wide association study (PheWAS) of Neanderthal- and Denisovan-introgressed alleles to characterize their contribution in EAS individuals and other ancestry groups available from the UK Biobank (UKB) ^38^. Our findings expand the understanding of how human evolutionary history influenced the genetic liability to complex traits, also providing evidence of the contribution of Denisovan introgression to physiological and pathological conditions in EAS populations.

## Results

### Partitioned Heritability Analysis

For the partitioned heritability analysis based on baseline and evolutionary annotations of the human genome, we identified a total of 37 and 39 traits with adequate SNP-*h*^*2*^ estimates (*z*-score ≥ 7) among those available in the BBJ (EAS participants) and the ones matched from the UKB (EUR participants), respectively. As expected, we observed a strong correlation between effective sample size and heritability *z*-score in both EAS and EUR (ρ = 0.75, p = 1.86 × 10^−13^ and ρ = 0.82, p = 2.20 × 10^−16^, respectively) (Supplemental Table 3). We identified several differences between EAS and EUR enrichments of genome structure and function annotations (Supplemental Table 4). The enrichment of three traits (i.e., blood sugar, mean corpuscular volume, non-albumin protein) was different for H3k27 active enhancer acetylation (H3K27ac) in EAS and EUR (most significant difference: non-albumin protein was more enriched for this functional annotation in EAS compared to EUR (EAS: 2.88-fold enrichment, p = 1.22 × 10^−18^, EUR: 1.11-fold enrichment, p = 0.080, EAS-EUR difference: p = 2.96 × 10^−12^)). Moreover, albumin/globulin ratio was depleted for H3K27ac flanking region in EAS (−6.14-fold depletion, p = 0.001), but it was significantly enriched in EUR (2.21-fold enrichment, p = 2.26 × 10^−10^; EAS-EUR difference: p = 2.72 × 10^−4^). The super-enhancer annotation was enriched in EAS (4.46-fold enrichment, p = 3.01 × 10^−17^), but not in EUR (1.14-fold enrichment, p = 0.105) with respect to non-albumin protein (EAS-EUR difference: p = 3.43 × 10^−12^). Background selection was more significantly enriched in lymphocyte count in EUR compared to EAS (EUR: 1.82-fold enrichment, p = 1.12 × 10^−18^; EAS: 1.30-fold enrichment, p = 2.18 × 10^−4^; EAS-EUR difference: p = 4.59 × 10^−4^). The enrichment of three traits (i.e., lymphocyte count, neutrophil count, non-albumin protein) was different for CpG content between EAS and EUR. The most significant difference was for non-albumin protein, which was more enriched for this functional annotation in EAS compared to EUR (EAS: 1.51-fold enrichment, p = 1.37 × 10^−11^, EUR: 1.09-fold enrichment, p = 2.36 × 10^−6^, EAS-EUR difference: 2.89 × 10^−6^).

We also observed several enrichments for evolutionary features in the SNP-*h*^*2*^ of traits and diseases assessed in EAS and EUR individuals (Table 1, Supplemental Table 5). In line with previous studies ^12,39^, the strongest enrichments in both ancestry groups were observed for annotations related to genic and LOF regions. In EAS, 89% and 68% of the traits analyzed in EAS had significant SNP-*h*^*2*^ enrichments for genic and LOF regions (Table 1, FDR q<0.05). Platelet count was the most significantly enriched trait in both genic and LOF regions (1.33-fold enrichment, p = 7.64 × 10^−12^ and 2-fold enrichment, p = 1.98 × 10^−8^, respectively). Due to the much larger sample size available, 100% of the traits analyzed showed significant SNP-*h*^*2*^ enrichments for these functional annotations in EUR^10^. Accordingly, we identified several significant enrichments related to B-statistic values in EUR (i.e., reduction in allelic diversity due to purifying selection) ^1^. Due to the much larger sample GWAS size, all phenotypes in EUR showed FDR significant enrichment in at least one of the B-statistic value thresholds. Similar to other studies conducted in EUR ^17,40^, we did not identify SNP-*h2* enrichment for positive selection signatures in our EAS and EUR analyses (Supplemental Table 5). With respect to archaic introgression, we identified one FDR-significant SNP-*h2* enrichment: Denisovan-introgressed loci for coronary artery disease in EAS (1.7-fold enrichment, p = 0.003).

**Table 1.** Enrichment of natural selection and functional annotation measures. *P* values of significant enrichments of genic, loss-of-function (LoF) intolerant and Denisovan-introgressed loci, and three genomic annotations of background selection.

| Trait | Annotation | East Asian |  |  | European |  |  | EAS-EUR Difference |  |
| --- | --- | --- | --- | --- | --- | --- | --- | --- | --- |
|  |  | Fold-enrichment | SE | P-value | Fold-enrichment | SE | P-value | Z score | P-value |
| Alanine aminotransferase | B top1 | 0.855 | 0.349 | 0.983 | 2.728 | 0.614 | 0.005 | -2.652 | 0.008 |
|  | B top2 | -13.622 | 9.421 | 0.958 | 2.159 | 0.366 | 0.001 | -1.674 | 0.094 |
|  | Genic | 1.237 | 0.081 | 0.002 | 1.315 | 0.034 | 2.48E-18 | -0.885 | 0.376 |
|  | LoF intolerant | 1.272 | 0.232 | 0.227 | 1.615 | 0.121 | 2.18E-07 | -1.311 | 0.190 |
| Albumin | B top05 | -1.971 | 2.266 | 0.718 | 3.950 | 1.484 | 0.049 | -2.186 | 0.029 |
|  | B top1 | 1.135 | 0.675 | 0.983 | 3.158 | 0.947 | 0.024 | -1.740 | 0.082 |
|  | B top2 | 8.223 | 7.181 | 0.958 | 3.734 | 0.670 | 6.11E-05 | 0.622 | 0.534 |
|  | Genic | 1.380 | 0.092 | 1.82E-05 | 1.297 | 0.027 | 2.97E-19 | 0.866 | 0.386 |
|  | LoF intolerant | 1.532 | 0.284 | 0.053 | 1.648 | 0.110 | 1.59E-09 | -0.381 | 0.704 |
| Albumin/globulin ratio | B top05 | 2.151 | 2.822 | 0.718 | 4.159 | 1.164 | 0.006 | -0.658 | 0.511 |
|  | B top1 | 1.597 | 0.453 | 0.849 | 2.719 | 0.597 | 0.004 | -1.498 | 0.134 |
|  | B top2 | 1.794 | 1.880 | 0.953 | 2.943 | 0.632 | 0.002 | -0.579 | 0.563 |
|  | Genic | 1.437 | 0.079 | 1.35E-09 | 1.310 | 0.030 | 2.07E-16 | 1.502 | 0.133 |
|  | LoF intolerant | 2.056 | 0.283 | 5.44E-05 | 1.661 | 0.117 | 3.37E-09 | 1.294 | 0.196 |
| Aspartate aminotransferase | B top05 | -1.507 | 1.256 | 0.718 | 3.323 | 1.120 | 0.033 | -2.870 | 0.004 |
|  | B top1 | 2.930 | 2.220 | 0.983 | 2.985 | 0.647 | 0.002 | -0.024 | 0.981 |
|  | B top2 | -12.232 | 6.942 | 0.958 | 2.742 | 0.496 | 0.000285 | -2.152 | 0.031 |
|  | Genic | 1.287 | 0.067 | 2.26E-06 | 1.314 | 0.030 | 2.39E-22 | -0.371 | 0.711 |
|  | LoF intolerant | 1.503 | 0.213 | 0.010 | 1.596 | 0.120 | 7.26E-08 | -0.381 | 0.703 |
| Asthma | B top05 | -2.439 | 3.102 | 0.950 | 2.329 | 0.629 | 0.034 | -1.506 | 0.132 |
|  | B top1 | 3.842 | 2.358 | 0.983 | 2.486 | 0.524 | 0.004 | 0.561 | 0.575 |
|  | B top2 | -8.241 | 1.204 | 0.961 | 2.096 | 0.459 | 0.016 | -8.020 | 1.06E-15 |
|  | Genic | 1.008 | 0.085 | 0.928 | 1.123 | 0.040 | 0.001 | -1.235 | 0.217 |
|  | LoF intolerant | 0.869 | 0.236 | 0.575 | 1.223 | 0.095 | 0.016 | -1.394 | 0.163 |
| Blood sugar | B top2 | -1.152 | 1.446 | 0.953 | 2.446 | 0.533 | 0.003 | -2.334 | 0.020 |
|  | Genic | 1.368 | 0.089 | 2.51E-05 | 1.293 | 0.065 | 2.09E-09 | 0.682 | 0.495 |
|  | LoF intolerant | 1.324 | 0.274 | 0.217 | 1.635 | 0.192 | 2.06E-04 | -0.930 | 0.352 |
| Blood urea nitrogen | Genic | 1.237 | 0.073 | 3.28E-04 | NA | NA | NA | NA | NA |
|  | LoF intolerant | 1.471 | 0.238 | 0.043 | NA | NA | NA | NA | NA |
| Body mass index | Genic | 1.167 | 0.065 | 0.007 | 1.125 | 0.017 | 7.18E-12 | 0.628 | 0.530 |
|  | LoF intolerant | 2.086 | 0.236 | 2.05E-07 | 1.682 | 0.069 | 1.46E-20 | 1.643 | 0.100 |
| Chloride | Genic | 1.361 | 0.105 | 3.80E-05 | NA | NA | NA | NA | NA |
|  | LoF intolerant | 1.722 | 0.338 | 0.019 | NA | NA | NA | NA | NA |
| Coronary artery disease | B top05 | 1.338 | 1.226 | 0.718 | 3.230 | 0.968 | 0.020 | -1.211 | 0.226 |
|  | B top2 | 1.051 | 1.058 | 0.953 | 2.400 | 0.473 | 0.002 | -1.164 | 0.244 |
|  | Denisovan | 1.691 | 0.877 | 0.003 | 2.149 | 1.522 | 0.449 | -0.261 | 0.794 |
|  | Genic | 1.232 | 0.047 | 5.47E-08 | 1.250 | 0.040 | 1.48E-09 | -0.282 | 0.778 |
|  | LoF intolerant | 1.575 | 0.181 | 3.36E-04 | 1.681 | 0.141 | 3.07E-07 | -0.460 | 0.645 |
| Diastolic blood pressure | B top05 | 2.871 | 2.759 | 0.718 | 2.907 | 0.913 | 0.039 | -0.012 | 0.990 |
|  | B top1 | 1.124 | 0.341 | 0.983 | 2.532 | 0.479 | 0.001 | -2.394 | 0.017 |
|  | B top2 | 3.116 | 1.191 | 0.958 | 2.323 | 0.298 | 9.33E-06 | 0.646 | 0.519 |
|  | Genic | 1.271 | 0.084 | 3.07E-04 | 1.230 | 0.024 | 5.69E-18 | 0.472 | 0.637 |
|  | LoF intolerant | 2.050 | 0.288 | 5E-05 | 1.751 | 0.093 | 4.46E-15 | 0.988 | 0.323 |
| Estimated glomerular filtration rate | B top05 | 1.234 | 1.484 | 0.718 | 4.724 | 1.209 | 0.002 | -1.823 | 0.068 |
|  | B top1 | 2.184 | 0.471 | 0.983 | 3.960 | 0.848 | 4.97E-04 | -1.830 | 0.067 |
|  | B top2 | 9.394 | 6.212 | 0.006 | 2.955 | 0.444 | 8.23E-06 | 1.034 | 0.301 |
|  | Genic | 1.277 | 0.050 | 1.33E-08 | 1.306 | 0.031 | 2.23E-21 | -0.483 | 0.629 |
|  | LoF intolerant | 1.931 | 0.203 | 3.96E-07 | 1.853 | 0.135 | 3.09E-09 | 0.320 | 0.749 |
| Hematocrit | Genic | 1.342 | 0.072 | 1.45E-07 | NA | NA | NA | NA | NA |
|  | LoF intolerant | 2.193 | 0.280 | 1.99E-06 | NA | NA | NA | NA | NA |
| Hemoglobin | Genic | 1.397 | 0.092 | 2.85E-07 | NA | NA | NA | NA | NA |
|  | LoF intolerant | 2.336 | 0.342 | 4.79E-06 | NA | NA | NA | NA | NA |
| Lymphocyte count | B top1 | 1.392 | 0.437 | 0.960 | 3.024 | 0.742 | 0.006 | -1.896 | 0.058 |
|  | B top2 | 2.193 | 1.216 | 0.293 | 2.722 | 0.484 | 3.32E-04 | -0.404 | 0.686 |
|  | Genic | 1.178 | 0.079 | 0.015 | 1.316 | 0.028 | 2.70E-20 | -1.636 | 0.102 |
|  | LoF intolerant | 1.523 | 0.246 | 0.024 | 1.671 | 0.110 | 5.54E-09 | -0.549 | 0.583 |
| Mean arterial pressure | B top05 | 1.536 | 2.312 | 0.950 | 2.550 | 0.766 | 0.045 | -0.416 | 0.677 |
|  | B top1 | 1.085 | 0.372 | 0.983 | 2.220 | 0.436 | 0.005 | -1.980 | 0.048 |
|  | B top2 | 1.833 | 1.321 | 0.958 | 2.083 | 0.280 | 7.84E-05 | -0.185 | 0.853 |
|  | Genic | 1.314 | 0.076 | 6.07E-06 | 1.219 | 0.024 | 2.03E-16 | 1.189 | 0.234 |
|  | LoF intolerant | 2.092 | 0.260 | 2.57E-06 | 1.784 | 0.096 | 7.51E-16 | 1.112 | 0.266 |
| Mean corpuscular hemoglobin | B top05 | 3.948 | 6.980 | 0.718 | 4.182 | 1.496 | 0.030 | -0.033 | 0.974 |
|  | B top1 | 2.015 | 0.836 | 0.849 | 3.802 | 1.055 | 0.007 | -1.327 | 0.184 |
|  | B top2 | 2.520 | 2.993 | 0.953 | 4.056 | 0.734 | 1.64E-05 | -0.498 | 0.618 |
|  | Genic | 1.303 | 0.075 | 8.83E-06 | 1.348 | 0.046 | 1.19E-12 | -0.515 | 0.607 |
|  | LoF intolerant | 1.723 | 0.278 | 0.006 | 1.705 | 0.167 | 6.71E-06 | 0.056 | 0.955 |
| Mean corpuscular volume | B top1 | 1.893 | 0.630 | 0.849 | 3.790 | 1.064 | 0.009 | -1.534 | 0.125 |
|  | B top2 | 3.423 | 3.146 | 0.953 | 3.715 | 0.666 | 4.42E-05 | -0.091 | 0.928 |
|  | Genic | 1.311 | 0.061 | 2.22E-07 | 3.715 | 0.666 | 4.42E-05 | -3.591 | 3.29E-04 |
|  | LoF intolerant | 1.657 | 0.243 | 0.006 | 1.699 | 0.154 | 4.14E-06 | -0.143 | 0.886 |
| Monocyte count | B top05 | 1.967 | 2.920 | 0.718 | 5.248 | 2.118 | 0.045 | -0.910 | 0.363 |
|  | B top1 | 2.456 | 1.347 | 0.849 | 3.793 | 1.004 | 0.005 | -0.796 | 0.426 |
|  | B top2 | 1.348 | 0.975 | 0.013 | 2.773 | 0.628 | 0.005 | -1.228 | 0.219 |
|  | Genic | 1.255 | 0.086 | 0.002 | 1.318 | 0.040 | 7.53E-13 | -0.660 | 0.509 |
|  | LoF intolerant | 1.548 | 0.240 | 0.018 | 1.557 | 0.150 | 8.76E-05 | -0.030 | 0.976 |
| Neutrophil count | B top05 | 1.800 | 1.885 | 0.950 | 4.148 | 1.106 | 0.004 | -1.074 | 0.283 |
|  | B top1 | 1.096 | 0.381 | 0.983 | 3.184 | 0.750 | 0.003 | -2.483 | 0.013 |
|  | B top2 | 4.964 | 0.986 | 0.958 | 2.751 | 0.518 | 0.001 | 1.985 | 0.047 |
|  | Genic | 1.235 | 0.080 | 0.002 | 1.256 | 0.025 | 3.62E-21 | -0.248 | 0.804 |
|  | LoF intolerant | 1.471 | 0.210 | 0.021 | 1.766 | 0.107 | 4.15E-11 | -1.255 | 0.210 |
| Non-albumin protein | B top05 | 4.440 | 5.075 | 0.718 | 4.291 | 1.014 | 0.001 | 0.029 | 0.977 |
|  | B top1 | 1.817 | 0.620 | 0.849 | 2.675 | 0.549 | 0.002 | -1.036 | 0.300 |
|  | B top2 | 2.184 | 2.456 | 0.953 | 2.847 | 0.540 | 0.001 | -0.264 | 0.792 |
|  | Genic | 1.468 | 0.094 | 2.72E-09 | 1.315 | 0.033 | 6.72E-16 | 1.532 | 0.125 |
|  | LoF intolerant | 2.373 | 0.301 | 9.41E-07 | 1.714 | 0.114 | 1.57E-10 | 2.043 | 0.041 |
| Platelet count | B top1 | 3.575 | 0.375 | 0.983 | 4.781 | 1.276 | 0.004 | -0.907 | 0.364 |
|  | B top2 | 2.644 | 0.641 | 0.958 | 3.744 | 0.586 | 6.10E-06 | -1.267 | 0.205 |
|  | Genic | 1.330 | 0.044 | 7.64E-12 | 1.366 | 0.026 | 7.71E-24 | -0.710 | 0.478 |
|  | LoF intolerant | 1.998 | 0.166 | 1.98E-08 | 1.984 | 0.142 | 5.76E-11 | 0.064 | 0.949 |
| Potassium | B top1 | 1.112 | 0.363 | 0.983 | 2.387 | 0.390 | 3.54E-04 | -2.393 | 0.017 |
|  | B top2 | 2.507 | 1.948 | 0.958 | 2.326 | 0.292 | 2.99E-06 | 0.092 | 0.927 |
|  | Genic | 1.306 | 0.078 | 1.36E-05 | 1.103 | 0.032 | 0.002 | 2.425 | 0.015 |
|  | LoF intolerant | 1.944 | 0.284 | 5.48E-05 | 1.641 | 0.115 | 7.51E-08 | 0.992 | 0.321 |
| Pulse pressure | Genic | 1.393 | 0.094 | 3.91E-07 | NA | NA | NA | NA | NA |
|  | LoF intolerant | 2.304 | 0.306 | 2.14E-06 | NA | NA | NA | NA | NA |
| Red blood cell count | B top05 | 1.639 | 2.885 | 0.718 | 4.790 | 1.325 | 0.004 | -0.993 | 0.321 |
|  | B top1 | 2.173 | 0.691 | 0.849 | 3.476 | 0.724 | 0.001 | -1.303 | 0.193 |
|  | B top2 | 5.614 | 1.771 | 0.958 | 3.033 | 0.487 | 1.83E-05 | 1.405 | 0.160 |
|  | Genic | 1.353 | 0.066 | 4.87E-08 | 1.130 | 0.025 | 4.54E-07 | 3.159 | 0.002 |
|  | LoF intolerant | 2.077 | 0.255 | 1.00E-05 | 1.561 | 0.092 | 1.19E-09 | 1.901 | 0.057 |
| Serum creatinine | B top05 | 8.840 | 5.950 | 0.021 | 4.724 | 1.209 | 0.002 | 0.678 | 0.498 |
|  | B top1 | 3.121 | 1.461 | 0.983 | 3.960 | 0.848 | 4.97E-04 | -0.497 | 0.619 |
|  | B top2 | 1.492 | 1.396 | 0.718 | 2.955 | 0.444 | 8.23E-06 | -0.998 | 0.318 |
|  | Genic | 1.289 | 0.052 | 7.98E-09 | 1.306 | 0.031 | 2.23E-21 | -0.277 | 0.782 |
|  | LoF intolerant | 1.865 | 0.194 | 9.85E-07 | 1.853 | 0.135 | 3.09E-09 | 0.050 | 0.960 |
| Smoking behaviors: Smoking initiation | Genic | 1.041 | 0.129 | 0.754 | NA | NA | NA | NA | NA |
|  | LoF intolerant | 1.379 | 0.324 | 0.248 | NA | NA | NA | NA | NA |
| Sodium | B top05 | 3.381 | 4.513 | 0.718 | 2.171 | 0.552 | 0.036 | 0.266 | 0.790 |
|  | B top1 | 1.362 | 0.715 | 0.983 | 1.953 | 0.329 | 0.003 | -0.750 | 0.453 |
|  | B top2 | -1.750 | 1.570 | 0.958 | 1.700 | 0.209 | 0.001 | -2.178 | 0.029 |
|  | Genic | 1.426 | 0.138 | 1.07E-04 | 1.130 | 0.025 | 4.54E-07 | 2.111 | 0.035 |
|  | LoF intolerant | 2.193 | 0.485 | 0.006 | 1.561 | 0.092 | 1.19E-09 | 1.283 | 0.200 |
| Systolic blood pressure | B top05 | 1.752 | 2.069 | 0.718 | 2.296 | 0.515 | 0.013 | -0.255 | 0.799 |
|  | B top1 | 1.014 | 0.389 | 0.985 | 2.035 | 0.334 | 0.002 | -1.991 | 0.047 |
|  | B top2 | 3.253 | 1.345 | 0.958 | 2.055 | 0.244 | 9.62E-06 | 0.876 | 0.381 |
|  | Genic | 1.341 | 0.070 | 8.58E-08 | 1.234 | 0.024 | 3.92E-18 | 1.439 | 0.150 |
|  | LoF intolerant | 2.143 | 0.241 | 8.95E-08 | 1.795 | 0.087 | 3.96E-18 | 1.356 | 0.175 |
| Total cholesterol | B top1 | 4.831 | 2.371 | 0.983 | 4.722 | 1.431 | 0.008 | 0.039 | 0.969 |
|  | B top2 | -7.322 | 9.190 | 0.953 | 2.873 | 0.781 | 0.012 | -1.105 | 0.269 |
|  | Genic | 1.359 | 0.086 | 2.71E-05 | 1.308 | 0.034 | 3.49E-13 | 0.553 | 0.580 |
|  | LoF intolerant | 1.287 | 0.303 | 0.308 | 1.432 | 0.174 | 0.011 | -0.414 | 0.679 |
| Total protein | B top05 | 5.388 | 5.538 | 0.718 | 4.096 | 0.826 | 2.83E-04 | 0.231 | 0.818 |
|  | B top1 | 1.956 | 0.514 | 0.849 | 2.736 | 0.520 | 0.001 | -1.066 | 0.286 |
|  | B top2 | 1.610 | 1.907 | 0.953 | 2.825 | 0.405 | 5.76E-06 | -0.623 | 0.533 |
|  | Genic | 1.395 | 0.093 | 2.96E-07 | 1.310 | 0.027 | 5.72E-18 | 0.874 | 0.382 |
|  | LoF intolerant | 2.111 | 0.278 | 2.98E-05 | 1.745 | 0.129 | 1.75E-10 | 1.197 | 0.231 |
| Type 2 diabetes | B top05 | 2.218 | 1.431 | 0.718 | 2.204 | 0.501 | 0.016 | 0.009 | 0.993 |
|  | B top1 | 1.085 | 0.292 | 0.983 | 2.109 | 0.392 | 0.005 | -2.092 | 0.036 |
|  | B top2 | 5.468 | 3.974 | 0.293 | 2.090 | 0.254 | 6.08E-06 | 0.848 | 0.396 |
|  | Genic | 1.171 | 0.034 | 8.13E-07 | 1.147 | 0.032 | 8.17E-06 | 0.523 | 0.601 |
|  | LoF intolerant | 1.955 | 0.179 | 1.56E-07 | 1.608 | 0.123 | 2.03E-06 | 1.596 | 0.110 |
| White blood cell count | B top05 | 5.024 | 3.190 | 0.718 | 5.029 | 1.350 | 0.003 | -0.001 | 0.999 |
|  | B top1 | 1.200 | 0.354 | 0.983 | 3.307 | 0.666 | 0.001 | -2.793 | 0.005 |
|  | B top2 | -5.583 | 7.351 | 0.953 | 2.757 | 0.479 | 2.79E-04 | -1.132 | 0.258 |
|  | Genic | 1.340 | 0.073 | 5.24E-07 | 1.253 | 0.023 | 3.31E-23 | 1.148 | 0.251 |
|  | LoF intolerant | 1.728 | 0.204 | 1.59E-04 | 1.730 | 0.100 | 8.96E-12 | -0.007 | 0.995 |

### Phenome-wide association study of Archaic introgressed loci

Although we observed only one SNP-*h2* enrichment for archaic introgression (i.e., Denisovan-introgressed loci for coronary artery disease in EAS), single loci inherited from Neanderthals and Denisovans can still contribute to the phenotypic variation of human populations ^20^. Therefore, we performed a PheWAS of archaic introgressed loci across multiple ancestry groups.

In EAS, we identified 45 LD-independent Denisovan-introgressed variants associated with 22 phenotypes (FDR q<0.05; Figure 1, Supplemental Table 6). These were related to 12 categories. The four most abundant categories were immunological phenotypes (eight LD-independent associations; most significant association for rheumatoid arthritis: rs59185462, beta = 0.822, p = 1.91 × 10^−105^ and chronic hepatitis B: rs115888238, beta = −0.618, p = 2.87 × 10^−18^), metabolic phenotypes (eight LD-independent associations; most significant association for type 2 diabetes: rs79748283, beta = −0.139, 5.07 × 10^−24^), cardiovascular phenotypes (seven LD-independent associations; most significant association for coronary artery disease: rs3784317, beta = −0.073, p = 5.97 × 10^−13^ and arrhythmia: rs4788667, beta = 0.094, p = 8.28 × 10^−12^), and endocrine phenotypes (three LD-independent associations; most significant association for Graves’ disease: rs79517313, beta = −0.477, 3.90 × 10^−18^). With respect to Neanderthal introgression, we did not observe any FDR significant association in EAS (Figure 1, Supplemental Table 7).

**Figure 1.**
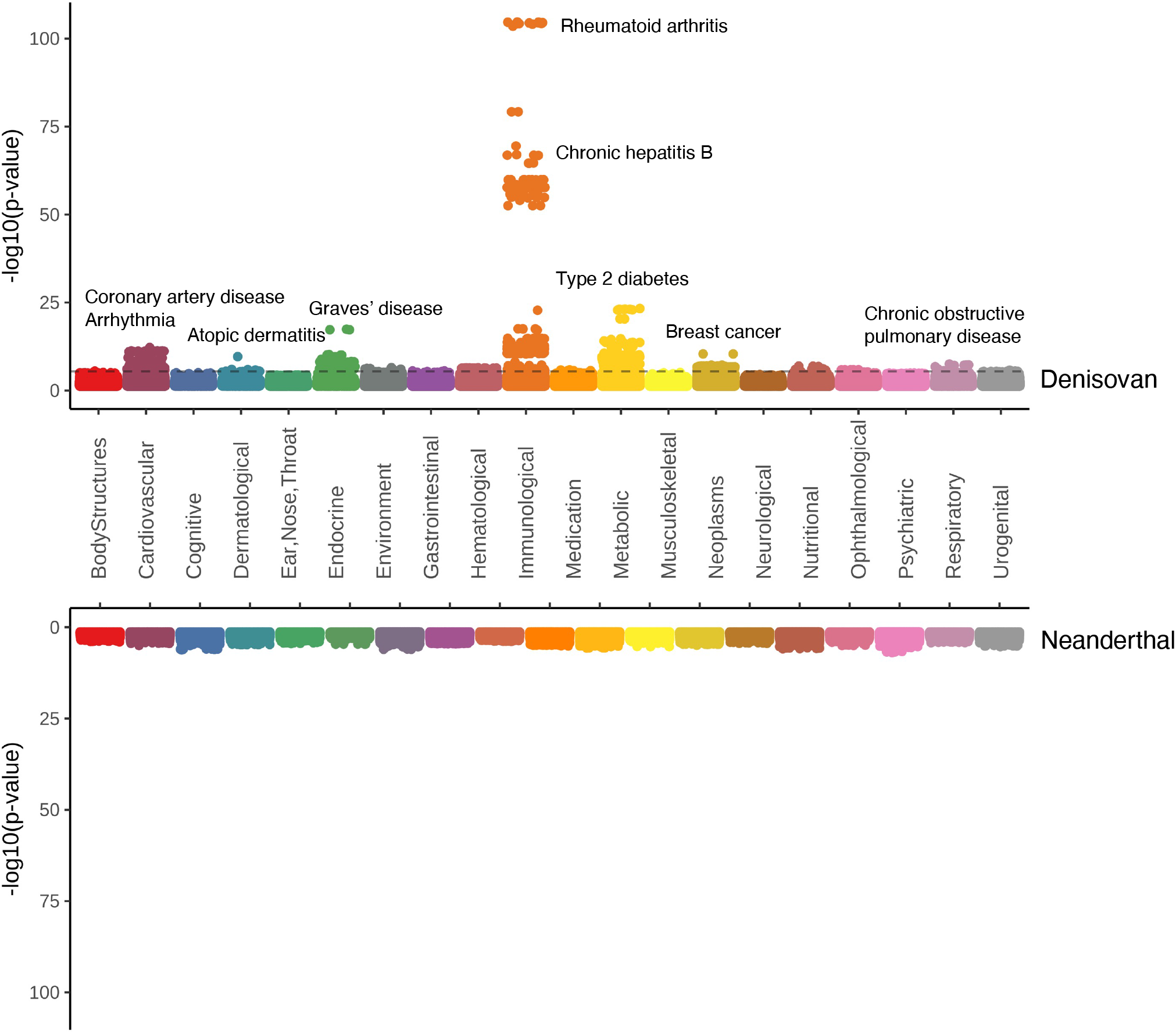
PheWAS Miami plot comparing Denisovan and Neanderthal introgressed variant associations with phenotypes in EAS. The −log10(p-values) for the Denisovan PheWAS are above the phenotype descriptions and those for the Neanderthal GWAS are below. The dashed line shows FDR-significant threshold (q < 0.05).

Because some of the Denisovan-introgressed variants also matched to the Neanderthal genome, we identified 347 LD-independent Denisovan-introgressed variants associated with 139 phenotypes in EUR (FDR q<0.05; Figure 2, Supplemental Table 8). These were related to all 20 phenotype categories, but the most abundant were hematological phenotypes (82 LD-independent associations, most significant association for *mean corpuscular hemoglobin*: rs17251430, beta = 0.158, p = 9.97 × 10^−304^) and metabolic traits (39 LD-independent association; most significant association for *Insulin-like growth factor 1*: rs17171240, beta = 0.211, p = 6.01 × 10^−85^ (Figure 2). In EUR, we identified 132 LD-independent Neanderthal-introgressed variants associated with 37 phenotypes (FDR q<0.05). These were related to 13 categories and the most abundant were metabolic phenotypes (82 LD-independent association, most significant association for *alkaline phosphatase*: rs11244089, beta = 0.096, p = 3.69 × 10^−116^) and musculoskeletal phenotypes (18 LD-independent association, most significant association for *contracture of palmar fascia*: rs117361340, beta = 0.678, p = 2.58 × 10^−46^) (Figure 2, Supplemental Table 9).

**Figure 2.**
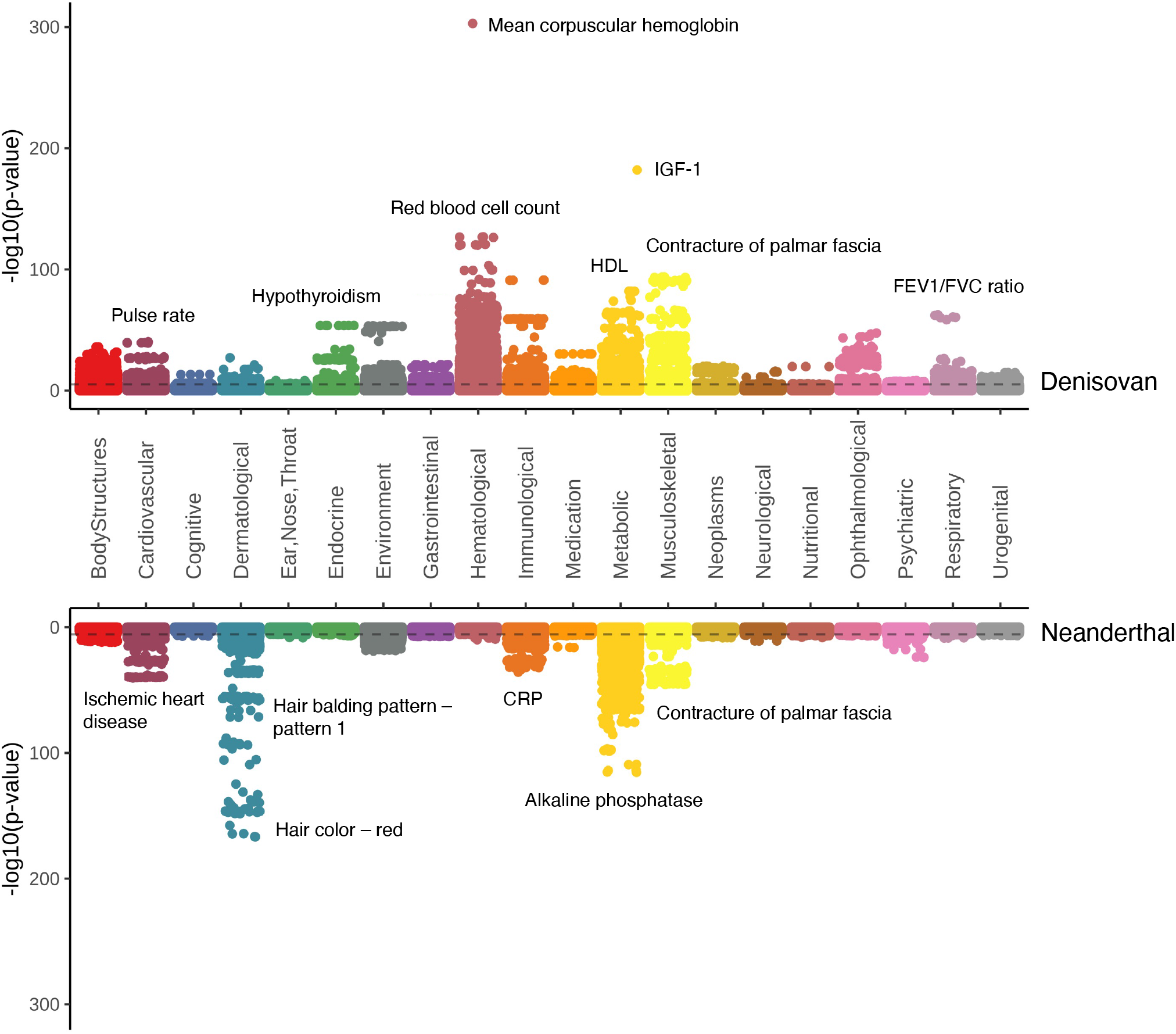
PheWAS Miami plot comparing Denisovan-and Neanderthal introgressed variant associations with phenotypes in EUR. The −log10(p-values) for the Denisovan PheWAS are above the phenotype descriptions and those for the Neanderthal GWAS are below. The dashed line shows FDR-significant threshold (q < 0.05).

Due to the limited sample size available, we did not find any FDR significant association for Denisovan or Neanderthal introgressed variants in CSA, AFR, MID, and AMR samples (Supplemental Tables 10-17).

### Over-representation test

Comparing the distribution of phenotypic classes associated with Denisovan-introgressed loci with the 7,221 phenotypes tested, we observed an over-representation for association with traits in the hematological category in EUR (2.99-fold enrichment, p = 5.89 × 10^−9^). Regarding phenotype classes associated with Neanderthal-introgressed loci, Metabolic category was overrepresented in EUR (1.70-fold enrichment, p = 1.70 × 10^−4^). No category was found to be overrepresented in the EAS analysis, which is likely due to the lower number of associations compared to EUR.

### Enrichment for biological processes, cellular components, and molecular functions

Considering the loci identified in our PheWAS, we tested the enrichment for biological processes, cellular components, and molecular functions. With respect to the Denisovan-or-Neanderthal introgressed loci identified in the EUR PheWAS, we identified 28 gene ontologies (FDR < 0.05) mostly related to cellular differentiation and epithelium development (Supplemental Table 18). Considering the Neanderthal loci identified in the PheWAS, we identified 30 gene-set enrichments (FDR < 5%) related to genomic regulation (Supplemental Table 19). Among them, we observed genes targeted by several microRNAs (miRNA, e.g. Hsa-miR-374b, FDR q = 9.27 × 10^−5^) and by different transcription factors (e.g., WT1 in human podocyte, FDR q = 9.27 × 10^−9^). Due to the limited number of loci identified in EAS PheWAS, no enrichment survived multiple testing correction in this ancestry group.

## Discussion

Modern populations present signatures of introgression from two archaic human species in their genomes ^28^. Our understanding of the phenotypic consequences of the alleles inherited from the admixture of anatomically modern humans with Denisovan and Neanderthal is still very limited. Previous studies showed that Neanderthal-introgressed loci are associated with immunological, neurological, psychiatric, metabolic, cardiovascular, and dermatological outcomes in EUR populations ^12,20,21,23^. In our study, we expanded this previous knowledge by testing for enrichment and depletion of SNP-*h*^*2*^ for loci related to Denisovan- and Neanderthal-introgression and several other evolutionary features across multiple traits in EAS and EUR populations. Additionally, we provide the first evidence of the consequences of Denisovan introgression across the human phenotypic spectrum in EAS and EUR. Among the results obtained from the present study, our findings highlight the specific contribution of Neanderthal and Denisovan introgression in the genetic liability to physiological and pathological conditions of EAS populations.

We found several associations for Denisovan-introgressed loci in EAS and EUR and Neanderthal-introgressed loci in EUR. Modern-day EUR populations do not carry or carry a low amount of Denisovan DNA ^36,37^, so we attribute the associations we found between EUR phenotypes and Denisovan-introgressed variants to the variants that match both Denisovan and Neanderthal genomes as they shared a common ancestor 600,000 years ago ^41^. In our evolution-focused SNP-*h*^*2*^ enrichment analysis, we detected an overabundance of genic and LoF intolerant loci in both EAS and EUR, suggesting that functionally important regions of the genome contribute to SNP-*h*^*2*^ to a different extent compared to the other annotations tested ^12,39,42,43^. Most of the traits tested were also enriched in CpG content, which is known to be positively correlated with genic content ^11^. Genic and LoF regions are strongly under negative selection ^10^. While most EUR phenotypes (76%) were highly enriched in B-statistic values, we only found one FDR-significant association in EAS (serum creatinine). A similar disparity between EUR and EAS findings was also present for the B-statistic continuous annotation. This is likely due to the much larger sample size available in EUR and may not reflect a general lack of evidence for background selection in EAS populations (Supplemental Table 3). We also observed that some functional enrichments were significantly more enriched in EAS than in EUR. For example, the super-enhancer annotation was enriched in EAS, but not in EUR. Genomic regions including enhancers have been shown to present an accelerated evolutionary rate, which is a signature of positive selection ^13^. However, similar to previous studies ^17,40^, none of the positive-selection annotations tested was significant in the two populations tested.

Leveraging genome-wide information in EAS, we observed that Denisovan-introgressed loci are more associated with the variation of a cardiovascular trait (coronary artery disease) than expected by chance. Two related cardiovascular phenotypes, myocardial infarction, and coronary atherosclerosis were previously associated with Neanderthal-introgressed loci in EUR ^20^. In our study, we have genome-wide and locus-level evidence that Denisovan introgression is linked to coronary artery disease: Denisovan-inherited variants explained a significantly higher proportion of coronary artery disease SNP-*h*^*2*^ and *ADAMTS7* rs3784317 and *CDKN2B-AS1* rs77953206 showed phenome-wide significant association with this trait. Furthermore, *ZFHX3* rs4788667 and rs78673846 showed phenome-wide significant association with arrhythmia. *ZFHX3* was identified in a previous atrial fibrillation GWAS ^44^. Our findings highlight that some of the pathogenic mechanisms involved in heart diseases in Denisovans could be shared with EAS populations. These EAS findings appear to be ancestry specific, because none of these associations were identified in the much larger EUR sample.

In our study, we found locus level evidence that Denisovan introgression is linked to type 2 diabetes in EAS: *KCNQ1* rs79748283, *AC104063*.*1* rs117233795, *LINC01153* rs11199817, *GLP1R* rs2268636, *C10orf67* rs79365475, and *PPIEL* rs61779359 showed phenome-wide significance with this trait. Among the loci identified in our PheWAS, prior research supports their association with type 2 diabetes, including a previous multi-ancestry association between T2D and *KCNQ1* ^45,46^, a genome-wide significant association between T2D and *LINC01153* ^47^, and the use of glucagon receptor agonists to treat certain types of T2D ^48,49^. Moreover, several missense mutations in its receptor were associated with insulin secretion and sensitivity impairment in Japanese patients ^50^. Our results suggest that both Denisovan and Neanderthal introgression may play a role in the development of type 2 diabetes in modern human populations. Most notably, we observed several other associations with Denisovan and Neanderthal alleles to support this notion. For instance, our association with insulin-like growth factor-1 (IGF-1) reinforces its utility as a biomarker for metabolic syndrome and T2D.^51^. Thus, this further supports our hypothesis that Denisovan and Neanderthal introgression may play a role in type 2 diabetes of modern populations. We also found several lipid-related metabolic factors associated with Denisovan- and Neanderthal-introgressed variants in both EUR and EAS. A previous study showed that variants shared between Neanderthals and modern humans are enriched in genes involved in lipid metabolism in EUR populations^52^. Our study expands this to EAS individuals. Among the Denisovan-introgressed loci identified, previous research supports the association of *PPIEL* with HDL cholesterol ^53^, and *ZPR1* with increased serum triglycerides and dyslipidemia ^54,55^.

Several of these metabolic phenotypes were also found to be associated with Neanderthal-introgressed variants (alkaline phosphatase: *SURF6* rs11244089, apolipoprotein A: *ALDH1A2* rs12900622, creatinine: *SLC7A9* rs57910615, *UBE2H* rs79808490, *TFDP2* rs73233892, cystatin C: *GRB10* rs73118816, glucose: *NOSTRIN* rs2433680, gamma glutamyltransferase: *SIGLEC1* rs12624921). Among them, *ALDH1A1* regulates adipogenesis, and can suppress *ALDH1A2* ^56^. *GRB10* expression was elevated in kidneys of diabetic mice ^57^, and *SLC7A9* was associated with chronic kidney disease ^58^. The convergent associations of Neanderthal and Denisovan-introgressed variants suggest that part of the genetic regulation of glucose and lipid metabolism and kidney function is inherited from archaic humans.

Actinic keratosis – a pre-cancerous skin condition – was previously associated with Neanderthal introgressed loci in EUR populations ^20^. Although actinic keratosis can be often observed in patients with atopic dermatitis, patients with atopic dermatitis do not appear to be at greater risk for developing the disease ^59^. Therefore, the locus-level evidence we found of *GLB1* rs76257456 associated with atopic dermatitis in our PheWAS can be considered an independent association. The association of this gene with atopic dermatitis was previously identified also in an independent Japanese cohort ^60^. While actinic keratosis is a dermatological condition caused by sun exposure, atopic dermatitis can be improved with sun exposure ^61^. Neanderthal introgression has been previously hypothesized to have a role in skin pigmentation and adaptation to ultraviolet radiation levels outside the African continent ^23,24^. Our findings highlight that Denisovan introgression may have played a similar role in EAS populations. Regarding Neanderthal-introgressed variants in EUR, red hair color and balding pattern were significantly associated with several variants (red hair color: *ANKRD11* rs60733936, *FANCA* rs11646374, *SLC24A4* rs77004437, *GALNS* rs75987792, *TPCN2* rs75840048, hair balding pattern: *LINC02210-CRHR1* rs62057113). Some of the variants were previously associated with hair color ^62–64^. A previous study concluded that Neanderthal variants contribute to light and dark tones of hair color, implying that Neanderthals were also variable in these traits ^21^. Although this previous investigation concluded that red hair is a uniquely human feature ^21^, we found that variants inherited from Neanderthals could also be associated with red hair. Therefore, more studies are needed to understand this possible association.

Neanderthal and Denisovan admixture have been previously associated with the diversity of innate immunity genes in modern humans ^20,65,66^. Neanderthal introgression was previously linked to immunological phenotypes, rheumatoid arthritis ^27^ and chronic hepatitis B ^67^, and Graves’ disease ^27^ in EUR and EAS. Denisovan introgression was linked to several immune processes, e.g., antiviral immune response, HIV-1 DNA integration, and cytokine signaling ^66^. In our study, we identified multiple Denisovan-introgressed loci associated with autoimmune disorders (rheumatoid arthritis and Graves’ disease) and viral diseases (chronic hepatitis B) in EAS. One of the loci identified (i.e., *HLA-B*) was identified as a specific risk factor for Graves’ disease in Asian populations ^68^. Our novel findings expand further our understanding of how human evolutionary history shaped the genetic regulation of immune function in worldwide populations.

In our study, chronic obstructive pulmonary disease (COPD) was associated with the *HYKK* rs79093205 Denisovan-introgressed variant in EAS. This phenotype has not been linked to Neanderthal introgression in EUR. However, it was found that a haplotype region on chromosome 3 inherited from Neanderthals was is a risk locus for respiratory failure upon COVID-19 infection ^69^. Several *HYKK* variants were previously associated with COPD in a GWAS in EUR and AFR ^70^. In the Denisovan PheWAS in EUR, we found that the *MEGF6* rs2096100 variant was related to another respiratory trait, the FEV1/FVC ratio. Because *HYKK* and *MEGF6* primary function is related to tissue development ^71,72^, we hypothesize that archaic-introgressed loci may contribute to pulmonary function independently from immune-related pathways.

Among novel phenotypes related to introgressed variants, pancreatic, and breast cancers were associated with two Denisovan-introgressed variants (rs12615584 and rs12143332, respectively). Previously, Neanderthal introgressed haplotypes were associated with prostate cancer ^27^. These convergent findings support that some molecular mechanisms linked to cancer susceptibility are shared between archaic humans and modern populations.

Considering the introgressed loci identified in our PheWAS analyses, we identified an overrepresentation of certain molecular mechanisms. These included processes related to cellular differentiation and development. In particular, many of them were related to kidney function. Dietary changes contributed to the human evolution of kidney ^73^. Since archaic introgression contributed to the adaptation of modern humans to different environments ^74^, we hypothesize that some of the adaptation to dietary changes may be due to introgressed loci regulating kidney function. We also observed that several Neanderthal-introgressed loci identified were related to transcriptomic regulation via transcription factors (i.e., proteins that control transcription from DNA to mRNA) and miRNA (i.e., non-coding RNA responsible for RNA silencing and post-transcriptional gene expression regulation). Previous studies showed that miRNA seed regions are under significant background selection ^75^ and that miRNA seed region can be affected by variants introgressed from Neanderthals ^76^. Moreover, Neanderthal-introgressed sequences in modern humans have a measurable impact on gene expression variation ^77^, and Neanderthals differed from humans rather in their regulatory sequences instead of protein-coding sequences ^78^.

In conclusion, our study expands the understanding of how evolutionary pressures shaped the genetic architecture of human traits and diseases across worldwide populations. The present findings highlight how certain evolutionary processes are shared among human groups while others may have had a specific contribution to certain populations. In particular, we present evidence that Neanderthal and Denisovan introgression contributed specifically to shape the genetics of complex traits in East Asia. This strongly supports the need to expand the representation of human diversity in genetic research to ensure a comprehensive understanding of the complex dynamics by which the variation in the human genome is linked to the variation in the human phenome.

## Methods

### Datasets

GWAS statistics were accessed from BBJ ^34,35^ and the UKB ^79^. BBJ is a registry of over 200,000 Japanese patients including information about 47 diseases and 59 quantitative traits (Supplemental Table 1) ^34,35^. The UKB dataset provides information regarding more than 7,000 phenotypes assessed in up to 500,000 participants from six ancestry groups ^38^. We obtained genome-wide association statistics from a pan-ancestry genetic analysis of the UKB (Pan-UKB). A detailed description of this analysis is available at https://pan.ukbb.broadinstitute.org. Briefly, multi-ancestry genome-wide association analyses of 7,221 phenotypes were performed using a generalized mixed model association testing framework. Ancestry-specific GWAS statistics are available for six genetically-determined ancestry groups: European (N = 420,531), Central/South Asian (CSA, N = 8,876), African (AFR, N= 6,636), East Asian (N = 2,709), Middle Eastern (MID, N = 1,599), Admixed American (AMR, N = 980).

### Linkage Disequilibrium Score Regression

The Linkage Disequilibrium Score Regression method (LDSC) was used to quantify the enrichment of evolutionary annotations in the SNP-*h*^*2*^ of each trait ^5^. For each binary trait, the effective sample size was calculated as recommended previously ^80^. The major histocompatibility complex region was excluded from the analysis due to its complex LD structure. SNP-*h*^*2*^ was calculated for each BBJ phenotype and, as recommended ^81^, those with an estimated SNP-*h*^*2*^ z score ≥ 7 were selected for the partitioned SNP-*h*^*2*^ analysis. To compare BBJ EAS participants with other ancestry groups, we selected 79 UKB traits that were assessed similarly to those available in BBJ. Due to the limited sample size in UKB for other ancestry groups, we limited our partitioned SNP-*h*^*2*^ analysis to the data derived from UKB EUR participants. Accordingly, we used LD scores generated from the 1000 Genome Project Phase 3 EAS and EUR reference panels to analyzes GWAS data generated from BBJ and UKB, respectively ^82^.

SNP-*h*^*2*^ partitioning^81^ was performed considering 95 baseline genomic annotations characterizing important molecular properties such as allele frequency distributions, conserved regions of the genome, and regulatory elements ^9^, 11 annotations (five binary annotations and corresponding flanking annotations and one continuous count annotation) ^83^, four human promoter annotations (promoter, promoter from the Exome Aggregation Consortium ^84^, genes, and two corresponding flanking annotations) ^85^, three human enhancer annotations (enhancer and corresponding flanking annotation + enhancer-enhancer conservation count) ^85^, two human promoter sequence age annotations (including one flanking annotation) ^86^, and two human enhancer sequence age annotation (including one flanking annotation) ^86^. We created additional genome-wide annotations for Denisovan ^87,88^ and Neanderthal ^28,87–89^-introgressed, positively selected ^14,17,90^, negatively selected ^1,91^, genic and LoF intolerant ^39^ positions using the publicly available datasets from the original publications. Denisovan or Neanderthal (hereinafter “Denisovan”) (N = 29,195) and Neanderthal-introgressed (N = 49,793) positions were derived from the Sprime dataset ^88^, which identified these archaic-introgressed positions from the 1000 Genome Project with respect to the Japanese population sample (i.e., Japanese in Tokyo, Japan). We selected this population to match the genetic diversity of the BBJ participants. We defined Denisovan SNPs as those matching the Denisovan genome (in some cases they also matched the Neanderthal genome, as approximately 20% of the Neanderthal-introgressed variants are also carried by Denisovans ^87^). We used this approach due to the low number of Denisovan-only variants in the Japanese population ^87^. Neanderthal SNPs we selected were matched to the Neanderthal genome. The contribution of Neanderthal ancestry was also assessed by another method that compares human (we used data specific for the Japanese population) and Neanderthal genomes, inferring the probability of admixture with Neanderthals for each human haplotype (Neanderthal local ancestry) ^28,89^. Positive selection was tested based on integrated haplotype score (iHS) for Asian populations, which reports detection of positive selection during the last ∼30,000 years based on the detection of abnormally long haplotypes ^16^. Cross-population extended haplotype homozygosity (XP-EHH) comparing East Asian and European ancestries based on 1000 Genomes was also used to detect differential selective pressure since the two populations diverged ^17^. The B-statistic for East Asians was used to assess background selection. B measures phylogenetic information from other primates to determine the reduction in allelic diversity in humans due to purifying selection ^1^. The ExAC database was used to annotate genic and LoF-intolerant regions of the genome. Each gene was assigned a probability of LoF intolerance (pLI) score ^39^. Continuous evolutionary measurements were analyzed as top 2%, top 1%, and top 0.5% of scores genome-wide as binary annotations as recommended before due to the difficulty of setting specific thresholds to define regions under negative- and positive selection ^12,42,91^. The evolutionary annotations used in EUR are reported in Wendt *et al* ^12^. Apart from those reported in this study, we created additional annotations for Denisovan- and Neanderthal-introgressed positions as explained before. We applied False Discovery Rate (FDR) multiple-testing correction (*q* ≤0.05) ^92^ accounting for the number of phenotypes tested. Partitioned SNP-*h*^*2*^ in LDSC analyzes a large linear model including all annotations simultaneously such that enrichment values for a single annotation reflect independence from all other annotations in the model.

### Phenome-wide association study

To increase the resolution of our investigation (from heritability enrichment to single-variant contribution), we conducted a PheWAS of Denisovan- and Neanderthal introgressed loci in EAS, EUR, and other ancestry groups available in the UKB (Admixed American, AMR; African, AFR; Middle Eastern, MID; Central-South Asian, CSA). PheWAS tests for association between a single variant and many phenotypes. To characterize further the contribution of SNPs with evidence of Denisovan and/or Neanderthal introgression, we investigated their association with >7,000 phenotypes in UKB and BBJ. The same Denisovan and Neanderthal-introgressed variants were used in the PheWAS than in the LDSC analysis (N = 29,295 and N = 49,793, respectively).

Our phenome-wide analysis included traits related to body structures, cardiovascular, cognitive, dermatological, ear-nose-throat, endocrine, environmental, gastrointestinal, hematological, immunological, medication, metabolic, musculoskeletal, neoplasms, neurological, nutritional, ophthalmological, psychiatric, respiratory, and urogenital domains (Supplemental Table 2) ^93^. We applied False Discovery Rate (FDR; *q* <0.05) ^92^ accounting for the number of phenotypes, variants, and ancestries tested to identify associations surviving multiple testing correction. Variants with minor allele frequency (MAF) ≤ 0.05 and the variants with the “low-confidence” flag (i.e., variants with alternate allele count in cases ≤ 3, alternate allele count in controls ≤ 3, or minor allele count (cases and controls combined) ≤20) in the Pan UKB analysis were excluded from the analysis. We performed LD clumping using PLINK 1.9 ^94^ with a r^2^=0.1 within 500 kb windows. The significant variants were annotated to genes using the SNP Nexus variant annotation tool ^95^.

### Gene Ontology Enrichment

The significant genes identified in each PheWAS were analyzed for gene ontology enrichment using the ShinyGO tool set ^96^ and functional and molecular annotations (e.g., molecular pathways and gene ontology) from Ensembl ^97^. We considered FDR q < 0.05 to identify enrichments surviving multiple testing correction.

### Over-representation test

To test for over-representation of certain phenotypic classes among the associations observed in the PheWAS, we calculated the significance of the phenotypic enrichment by a hypergeometric distribution test (https://systems.crump.ucla.edu/hypergeometric/) where *k* is the number of phenotypes with at least one LD-independent association within the phenotype category of interest, *s* is the number of phenotypes with at least one LD-independent association, *M* is the number of phenotypes within the phenotype category of interest, and *N* is the number of phenotypes tested.

## Acknowledgements

The authors acknowledge support from the Horizon 2020 Marie Sklodowska-Curie Individual Fellowship from the European Commission (101028810) and the National Institutes of Health (R33 DA047527, R21 DC018098, and F32 MH122058). We thank the participants and the investigators involved in the UK Biobank, Biobank Japan, and the Pan-UK Biobank analysis for making their data publicly available.

## Author Contributions

Designed research: DK, FRW, RP; Analyzed data: DK; Interpreted the results: DK, FRW, GAP, ADL, ST, RP; Wrote the initial draft of the manuscript: DK; Critically revised manuscript: FRW, GP, ADL, FDA, BCM, ST, RP. All authors approved the final draft.

## Data availability

Biobank Japan summary statistics: http://jenger.riken.jp/en/result. Pan-UK Biobank summary statistics: https://pan.ukbb.broadinstitute.org/downloads. Baseline genomic annotations: https://alkesgroup.broadinstitute.org/LDSCORE/. Integrated haplotype score (iHS): http://hgdp.uchicago.edu/Browser_tracks/iHS/. Cross-population extended haplotype homozygosity (XP-EHH): http://hgdp.uchicago.edu/Browser_tracks/XPEHH/. B-statistic: https://github.com/gmcvicker/bkgd/tree/7ae49926008406bfcc81aec419e5d314390338e1. Denisovan and Neanderthal positions: https://data.mendeley.com/datasets/y7hyt83vxr/1. Neanderthal local ancestry: https://reich.hms.harvard.edu/datasets/landscape-neandertal-ancestry-present-day-humans. ExAC database: https://gnomad.broadinstitute.org/.

## References

1. McVicker, G., Gordon, D., Davis, C. & Green, P. Widespread genomic signatures of natural selection in hominid evolution. PLoS Genet 5, e1000471 (2009).

2. International Schizophrenia Consortium et al. Common polygenic variation contributes to risk of schizophrenia and bipolar disorder. Nature 460, 748–752 (2009).

3. Yang, J. et al.. Common SNPs explain a large proportion of the heritability for human height. Nat Genet 42, 565–569 (2010).

4. Diabetes Genetics Replication and Meta-analysis Consortium et al. Bayesian inference analyses of the polygenic architecture of rheumatoid arthritis. Nat Genet 44, 483–489 (2012).

5. Bulik-Sullivan, B. K. et al.. LD Score regression distinguishes confounding from polygenicity in genome-wide association studies. Nat Genet 47, 291–295 (2015).

6. Finucane, H. K. et al.. Partitioning heritability by functional annotation using genome-wide association summary statistics. Nat Genet 47, 1228–1235 (2015).

7. Zeng, J. et al.. Signatures of negative selection in the genetic architecture of human complex traits. Nat Genet 50, 746–753 (2018).

8. Zeng, J. et al.. Widespread signatures of natural selection across human complex traits and functional genomic categories. Nat Commun 12, 1164 (2021).

9. Gazal, S. et al.. Linkage disequilibrium-dependent architecture of human complex traits shows action of negative selection. Nat Genet 49, 1421–1427 (2017).

10. O’Connor, L. J. et al.. Extreme Polygenicity of Complex Traits Is Explained by Negative Selection. The American Journal of Human Genetics 105, 456–476 (2019).

11. Phung, T. N., Huber, C. D. & Lohmueller, K. E. Determining the Effect of Natural Selection on Linked Neutral Divergence across Species. PLoS Genet 12, e1006199 (2016).

12. Wendt, F. R. et al.. Characterizing the effect of background selection on the polygenicity of brain-related traits. Genomics 113, 111–119 (2021).

13. Moon, J. M., Capra, J. A., Abbot, P. & Rokas, A. Signatures of Recent Positive Selection in Enhancers Across 41 Human Tissues. G3 (Bethesda) 9, 2761–2774 (2019).

14. Grossman, S. R. et al.. A composite of multiple signals distinguishes causal variants in regions of positive selection. Science 327, 883–886 (2010).

15. Weigand, H. & Leese, F. Detecting signatures of positive selection in non-model species using genomic data. Zoological Journal of the Linnean Society 184, 528–583 (2018).

16. Voight, B. F., Kudaravalli, S., Wen, X. & Pritchard, J. K. A map of recent positive selection in the human genome. PLoS Biol 4, e72 (2006).

17. Sabeti, P. C. et al.. Genome-wide detection and characterization of positive selection in human populations. Nature 449, 913–918 (2007).

18. Meyer, M. et al.. A High-Coverage Genome Sequence from an Archaic Denisovan Individual. Science 338, 222–226 (2012).

19. Prüfer, K. et al.. The complete genome sequence of a Neanderthal from the Altai Mountains. Nature 505, 43–49 (2014).

20. Simonti, C. N. et al.. The phenotypic legacy of admixture between modern humans and Neandertals. Science 351, 737–741 (2016).

21. Dannemann, M. & Kelso, J. The Contribution of Neanderthals to Phenotypic Variation in Modern Humans. The American Journal of Human Genetics 101, 578–589 (2017).

22. McArthur, E., Rinker, D. & Capra, J. A. Quantifying the contribution of Neanderthal introgression to the heritability of complex traits. http://biorxiv.org/lookup/doi/10.1101/2020.06.08.140087 (2020) doi:10.1101/2020.06.08.140087.

23. Sankararaman, S. et al.. The genomic landscape of Neanderthal ancestry in present-day humans. Nature 507, 354–357 (2014).

24. Vernot, B. & Akey, J. M. Resurrecting Surviving Neandertal Lineages from Modern Human Genomes. Science 343, 1017–1021 (2014).

25. Vernot, B. et al.. Excavating Neandertal and Denisovan DNA from the genomes of Melanesian individuals. Science 352, 235–239 (2016).

26. Skov, L. et al.. The nature of Neanderthal introgression revealed by 27,566 Icelandic genomes. Nature 582, 78–83 (2020).

27. Dannemann, M. The Population-Specific Impact of Neandertal Introgression on Human Disease. Genome Biology and Evolution 13, evaa250 (2021).

28. Sankararaman, S., Mallick, S., Patterson, N. & Reich, D. The Combined Landscape of Denisovan and Neanderthal Ancestry in Present-Day Humans. Curr Biol 26, 1241–1247 (2016).

29. Gittelman, R. M. et al.. Archaic Hominin Admixture Facilitated Adaptation to Out-of-Africa Environments. Current Biology 26, 3375–3382 (2016).

30. Choin, J. et al.. Genomic insights into population history and biological adaptation in Oceania. Nature 592, 583–589 (2021).

31. Taskent, O., Lin, Y. L., Patramanis, I., Pavlidis, P. & Gokcumen, O. Analysis of Haplotypic Variation and Deletion Polymorphisms Point to Multiple Archaic Introgression Events, Including from Altai Neanderthal Lineage. Genetics 215, 497–509 (2020).

32. Huerta-Sánchez, E. et al.. Altitude adaptation in Tibetans caused by introgression of Denisovan-like DNA. Nature 512, 194–197 (2014).

33. Teixeira, J. C. et al.. Widespread Denisovan ancestry in Island Southeast Asia but no evidence of substantial super-archaic hominin admixture. Nat Ecol Evol 5, 616–624 (2021).

34. Kanai, M. et al.. Genetic analysis of quantitative traits in the Japanese population links cell types to complex human diseases. Nat Genet 50, 390–400 (2018).

35. Ishigaki, K. et al.. Large-scale genome-wide association study in a Japanese population identifies novel susceptibility loci across different diseases. Nat Genet 52, 669–679 (2020).

36. Chen, L., Wolf, A. B., Fu, W., Li, L. & Akey, J. M. Identifying and Interpreting Apparent Neanderthal Ancestry in African Individuals. Cell 180, 677-687.e16 (2020).

37. Hubisz, M. J., Williams, A. L. & Siepel, A. Mapping gene flow between ancient hominins through demography-aware inference of the ancestral recombination graph. PLoS Genet 16, e1008895 (2020).

38. Bycroft, C. et al.. The UK Biobank resource with deep phenotyping and genomic data. Nature 562, 203–209 (2018).

39. Lek, M. et al.. Analysis of protein-coding genetic variation in 60,706 humans. Nature 536, 285–291 (2016).

40. Yao, Y. et al.. No Evidence for Widespread Positive Selection Signatures in Common Risk Alleles Associated with Schizophrenia. Schizophrenia Bulletin 46, 603–611 (2020).

41. Rogers, A. R., Bohlender, R. J. & Huff, C. D. Early history of Neanderthals and Denisovans. Proc Natl Acad Sci USA 114, 9859–9863 (2017).

42. Pardiñas, A. F. et al.. Common schizophrenia alleles are enriched in mutation-intolerant genes and in regions under strong background selection. Nat Genet 50, 381–389 (2018).

43. McArthur, E., Rinker, D. C. & Capra, J. A. Quantifying the contribution of Neanderthal introgression to the heritability of complex traits. Nat Commun 12, 4481 (2021).

44. Husser, D. et al.. Association of atrial fibrillation susceptibility genes, atrial fibrillation phenotypes and response to catheter ablation: a gene-based analysis of GWAS data. J Transl Med 15, 71 (2017).

45. Been, L. F. et al.. Variants in KCNQ1 increase type II diabetes susceptibility in South Asians: a study of 3,310 subjects from India and the US. BMC Med Genet 12, 18 (2011).

46. Unoki, H. et al.. SNPs in KCNQ1 are associated with susceptibility to type 2 diabetes in East Asian and European populations. Nat Genet 40, 1098–1102 (2008).

47. Vujkovic, M. et al.. Discovery of 318 new risk loci for type 2 diabetes and related vascular outcomes among 1.4 million participants in a multi-ancestry meta-analysis. Nat Genet 52, 680–691 (2020).

48. Simó, R. & Hernández, C. GLP-1R as a Target for the Treatment of Diabetic Retinopathy: Friend or Foe? Diabetes 66, 1453–1460 (2017).

49. Hegedüs, L., Moses, A. C., Zdravkovic, M., Le Thi, T. & Daniels, G. H. GLP-1 and Calcitonin Concentration in Humans: Lack of Evidence of Calcitonin Release from Sequential Screening in over 5000 Subjects with Type 2 Diabetes or Nondiabetic Obese Subjects Treated with the Human GLP-1 Analog, Liraglutide. The Journal of Clinical Endocrinology & Metabolism 96, 853–860 (2011).

50. Tokuyama, Y. et al.. Five missense mutations in glucagon-like peptide 1 receptor gene in Japanese population. Diabetes Res Clin Pract 66, 63–69 (2004).

51. Aguirre, G. A., De Ita, J. R., de la Garza, R. G. & Castilla-Cortazar, I. Insulin-like growth factor-1 deficiency and metabolic syndrome. J Transl Med 14, 3 (2016).

52. Khrameeva, E. E. et al.. Neanderthal ancestry drives evolution of lipid catabolism in contemporary Europeans. Nat Commun 5, 3584 (2014).

53. Wiltshire, S. A. et al.. Genetic control of high density lipoprotein-cholesterol in AcB/BcA recombinant congenic strains of mice. Physiol Genomics 44, 843–852 (2012).

54. Ueyama, C. et al.. Association of FURIN and ZPR1 polymorphisms with metabolic syndrome. Biomed Rep 3, 641–647 (2015).

55. Bai, W. et al.. Functional polymorphisms of the APOA1/C3/A4/A5-ZPR1-BUD13 gene cluster are associated with dyslipidemia in a sex-specific pattern. PeerJ 6, e6175 (2019).

56. Petrosino, J. M., Disilvestro, D. & Ziouzenkova, O. Aldehyde dehydrogenase 1A1: friend or foe to female metabolism? Nutrients 6, 950–973 (2014).

57. Yang, S. et al.. Amelioration of Diabetic Mouse Nephropathy by Catalpol Correlates with Down-Regulation of Grb10 Expression and Activation of Insulin-Like Growth Factor 1 / Insulin-Like Growth Factor 1 Receptor Signaling. PLoS One 11, e0151857 (2016).

58. Corredor, Z. et al.. Genetic Variants Associated with Chronic Kidney Disease in a Spanish Population. Sci Rep 10, 144 (2020).

59. Hajdarbegovic, E. et al.. Atopic dermatitis is not associated with actinic keratosis: cross□sectional results from the Rotterdam study. Br J Dermatol 175, 89–94 (2016).

60. Hirota, T. et al.. Genome-wide association study identifies eight new susceptibility loci for atopic dermatitis in the Japanese population. Nat Genet 44, 1222–1226 (2012).

61. Patrizi, A., Raone, B. & Ravaioli, G. M. Management of atopic dermatitis: safety and efficacy of phototherapy. CCID 511 (2015) doi:10.2147/CCID.S87987.

62. Söchtig, J. et al.. Exploration of SNP variants affecting hair colour prediction in Europeans. Int J Legal Med 129, 963–975 (2015).

63. Han, J. et al.. A genome-wide association study identifies novel alleles associated with hair color and skin pigmentation. PLoS Genet 4, e1000074 (2008).

64. Sitek, A. et al.. Selected gene polymorphisms effect on skin and hair pigmentation in Polish children at the prepubertal age. Anthropol Anz 73, 283–293 (2016).

65. Quach, H. et al.. Genetic Adaptation and Neandertal Admixture Shaped the Immune System of Human Populations. Cell 167, 643-656.e17 (2016).

66. Almarri, M. A. et al.. Population Structure, Stratification, and Introgression of Human Structural Variation. Cell 182, 189-199.e15 (2020).

67. Datta, S. Excavating new facts from ancient Hepatitis B virus sequences. Virology 549, 89–99 (2020).

68. Li, Y. et al.. Association between HLA-B*46 Allele and Graves Disease in Asian Populations: A Meta-Analysis. Int. J. Med. Sci. 10, 164–170 (2013).

69. Zeberg, H. & Pääbo, S. The major genetic risk factor for severe COVID-19 is inherited from Neanderthals. Nature 587, 610–612 (2020).

70. Lutz, S. M. et al.. A genome-wide association study identifies risk loci for spirometric measures among smokers of European and African ancestry. BMC Genet 16, 138 (2015).

71. Veiga-da-Cunha, M., Hadi, F., Balligand, T., Stroobant, V. & Van Schaftingen, E. Molecular identification of hydroxylysine kinase and of ammoniophospholyases acting on 5-phosphohydroxy-L-lysine and phosphoethanolamine. J Biol Chem 287, 7246–7255 (2012).

72. Wang, Y., Song, H., Wang, W. & Zhang, Z. Generation and characterization of Megf6 null and Cre knock-in alleles. Genesis 57, e23262 (2019).

73. Chevalier, R. L. Evolutionary Nephrology. Kidney Int Rep 2, 302–317 (2017).

74. Rotival, M. & Quintana-Murci, L. Functional consequences of archaic introgression and their impact on fitness. Genome Biol 21, 3 (2020).

75. Quach, H. et al.. Signatures of purifying and local positive selection in human miRNAs. Am J Hum Genet 84, 316–327 (2009).

76. Lopez-Valenzuela, M. et al.. An Ancestral miR-1304 Allele Present in Neanderthals Regulates Genes Involved in Enamel Formation and Could Explain Dental Differences with Modern Humans. Molecular Biology and Evolution 29, 1797–1806 (2012).

77. McCoy, R. C., Wakefield, J. & Akey, J. M. Impacts of Neanderthal-Introgressed Sequences on the Landscape of Human Gene Expression. Cell 168, 916-927.e12 (2017).

78. Petr, M., Pääbo, S., Kelso, J. & Vernot, B. Limits of long-term selection against Neandertal introgression. Proc Natl Acad Sci USA 116, 1639–1644 (2019).

79. Pan-UKB team (2020). https://pan.ukbb.broadinstitute.org.

80. Willer, C. J., Li, Y. & Abecasis, G. R. METAL: fast and efficient meta-analysis of genomewide association scans. Bioinformatics 26, 2190–2191 (2010).

81. Finucane, H. K. et al.. Partitioning heritability by functional annotation using genome-wide association summary statistics. Nat. Genet. 47, 1228–1235 (2015).

82. 1000 Genomes Project Consortium et al. A global reference for human genetic variation. Nature 526, 68–74 (2015).

83. Hujoel, M. L. A., Gazal, S., Hormozdiari, F., van de Geijn, B. & Price, A. L. Disease Heritability Enrichment of Regulatory Elements Is Concentrated in Elements with Ancient Sequence Age and Conserved Function across Species. Am J Hum Genet 104, 611–624 (2019).

84. Karczewski, K. J. et al.. The ExAC browser: displaying reference data information from over 60 000 exomes. Nucleic Acids Res 45, D840–D845 (2017).

85. Villar, D. et al.. Enhancer evolution across 20 mammalian species. Cell 160, 554–566 (2015).

86. Marnetto, D. et al.. Evolutionary Rewiring of Human Regulatory Networks by Waves of Genome Expansion. Am J Hum Genet 102, 207–218 (2018).

87. Browning, S. R., Browning, B. L., Zhou, Y., Tucci, S. & Akey, J. M. Analysis of Human Sequence Data Reveals Two Pulses of Archaic Denisovan Admixture. Cell 173, 53-61.e9 (2018).

88. Browning, S. Sprime results for 1000 Genomes non-African populations and SGDP Papuans. (2018) doi:10.17632/Y7HYT83VXR.1.

89. Durvasula, A. & Sankararaman, S. A statistical model for reference-free inference of archaic local ancestry. PLoS Genet 15, e1008175 (2019).

90. Grossman, S. R. et al.. Identifying recent adaptations in large-scale genomic data. Cell 152, 703–713 (2013).

91. Huber, C. D., DeGiorgio, M., Hellmann, I. & Nielsen, R. Detecting recent selective sweeps while controlling for mutation rate and background selection. Mol Ecol 25, 142–156 (2016).

92. Benjamini, Y. & Hochberg, Y. Controlling the False Discovery Rate: A Practical and Powerful Approach to Multiple Testing. Journal of the Royal Statistical Society: Series B (Methodological) 57, 289–300 (1995).

93. Watanabe, K. et al.. A global overview of pleiotropy and genetic architecture in complex traits. Nat Genet 51, 1339–1348 (2019).

94. Chang, C. C. et al.. Second-generation PLINK: rising to the challenge of larger and richer datasets. Gigascience 4, 7 (2015).

95. Oscanoa, J. et al.. SNPnexus: a web server for functional annotation of human genome sequence variation (2020 update). Nucleic Acids Research 48, W185–W192 (2020).

96. Ge, S. X., Jung, D. & Yao, R. ShinyGO: a graphical gene-set enrichment tool for animals and plants. Bioinformatics 36, 2628–2629 (2020).

97. Aken, B. L. et al.. Ensembl 2017. Nucleic Acids Res 45, D635–D642 (2017).

